# Dietary Fiber Modulates Macrophage Activity in a Microfluidic Model of Colonocyte-Microbiota Interactions in Colorectal Cancer

**DOI:** 10.1101/2024.10.23.619945

**Authors:** Daniel Penarete-Acosta, Mohet Mittal, Sanjukta Chakraborty, Arum Han, Arul Jayaraman

## Abstract

Dietary fiber has been consistently associated with a decreased risk of colorectal cancer (CRC) development. While the apoptotic effect of dietary fiber microbial fermentation products on tumor colonocytes is well established, the role of these products on other components of the tumor microenvironment remains unexplored. Tumor associated macrophages play a critical role in tumor development in the colon; however, the effect of dietary fiber fermentation by microbiota on macrophage-colonocyte interaction in colorectal cancer has been difficult to dissect due to a lack of complex *in vitro* models of CRC containing both immune cells and microbiota. Recently, we developed a microfluidic model that facilitates the coculture of CRC spheroids with complex microbial communities. Here, we expand our model to include macrophages and employ it to study the impact of dietary fiber on macrophage-colonocyte interaction. We optimized monocyte differentiation parameters *in vitro* and demonstrated the capacity of our model to recapitulate changes in microbiota composition and metabolic output associated with dietary fiber administration *in vivo*. Combinatorial coculture of colonocytes with microbiota and macrophages revealed that alterations in microbial production of SCFA derived from dietary fiber fermentation correlated with enhanced colonocyte death, possibly mediated by an increase in transcription of tumor pro-apoptotic signals by macrophages. Our work highlights the capacity of complex *in vitro* systems to study the role of microbial metabolism of dietary molecules on CRC colonocyte viability and macrophage activity.

## Introduction

Dietary fiber consumption has been extensively associated with lower risk of CRC in humans^1^. A causative link between consumption dietary fiber in the form of inulin and decreased incidence of aberrant crypt foci and tumor formation in mice has also been reported^2^. Mechanistically, dietary fiber is metabolized by the colonic microbiota, which increases the abundance of fermentative bacteria and the production of short-chain fatty acids (SCFA) such as butyrate, propionate, and acetate^3–5^. The capacity of these metabolites to induce apoptosis in colonocyte via mechanisms such as histone deacetylase inhibition is well established^6–11^. For these reasons, prebiotic intervention with dietary fiber has been proposed as a preventive strategy against CRC^12^.

The colorectal tumor microenvironment contains multiple types of immune cells that significantly impact tumor development and progression^13^. Tumor-associated macrophages (TAMs) are abundant in carcinomas^14,15^, where they exhibit either tumor supportive activity by favoring cancer cell proliferation^16,17^ and migration^18–20^, or tumor-suppressive activity by inducing tumor cell apoptosis via signals such as Tumor Necrosis Factor α (TNF-α)^21^ and TRAIL^22^. The activity of TAM is influenced by host-derived molecules present in the tumor microenvironment such as cytokines, chemokines, non-coding RNA, and oncoproteins^23,24^.

In the colon, the microbiota and its metabolites have also been shown to modulate the activity of macrophages. Experiments in germ-free mice have shown that the microbiota regulates macrophage recruitment and replenishment after the onset of inflammation, possibly via induction of chemokine production in colonocytes^25^. Several pathogens associated with CRC, including *Fusobacterium nucleatum*, *Streptococcus gallolyticus*, and *Enterococcus faecalis*, promote a proinflammatory and immunosuppressive microenvironment by targeting macrophages^26^. In terms of bacterial metabolites, lipopolysaccharide is a potent inducer of proinflammatory polarization in macrophages^27^, while the SCFA butyrate favors anti-inflammatory polarization in macrophages while boosting their phagocytic activity^28,29^. Despite the high abundance of TAMs in tumors and the modulation of macrophage activity by microbiota, the effect of dietary fiber-induced changes in microbiota composition and function on TAM activity in CRC is not fully understood.

Previously, we developed a microfluidic device to study the interaction between colonocyte spheroids and colonic microbiota^30^. Here, we expand the physiological relevance of this model by co-culturing macrophages with colonocytes in spheroids. We hypothesize that inulin fermentation by colonic microbiota impacts macrophage-colonocyte interaction. We use metabolomics, metagenomics, and gene expression analysis to dissect the tripartite interaction between inulin, microbiota, and macrophages in our expanded CRC tumor microenvironment model. Our results contribute to our understanding of the effect of diet on host-microbiota interactions in the context of colorectal cancer and demonstrate the usefulness of physiologically relevant *in vitro* models to study these interactions.

## Results

### Inclusion of macrophages in an in vitro coculture model of CRC colonocyte-microbiota

To expand the physiological relevance of our previously developed colonocyte-microbiota coculture model^30^, we incorporated macrophages derived from the monocyte cell line THP-1. It is well-established that *in vitro* differentiation of THP-1 cells with PMA results in a macrophage-like phenotype, but the PMA concentration and treatment time widely vary among studies and significantly impact differentiation success^31–33^. Therefore, we optimized PMA concentration and treatment time to maximize the development of macrophages and the expression of the macrophage surface marker Cluster of Differentiation 11b (CD11b). THP-1 monocytes in culture are globular and remain in suspension. Treatment of THP-1 cells with PMA resulted in cell attachment to the bottom of the culture plate with changes in cell size and morphology (**Figure 1A**). After PMA treatment for 24 h, the attached cells were circular in shape and displayed increased granularity under phase-contrast microscopy. As the treatment time increased beyond 48 h, cells became elongated at PMA concentrations greater than 50 ng/mL, while cells treated with lower concentrations remained globular and loosely attached. CD11b expression increased with treatment time and was maximum after 72 h of treatment with a PMA concentration of 50 ng/mL, as assessed by flow cytometry. Minimal differences in cell morphology were observed with higher PMA concentrations for the same period of time (**Figure 1A**). Under these conditions, the mean fluorescence intensity in PMA-treated cells increased 75% compared to untreated cells (**Figure 1B, C**). Therefore, treatment with a PMA concentration of 50 ng/mL for 72 h was selected to induce THP-1 differentiation in device culture experiments.

**Figure 1.**
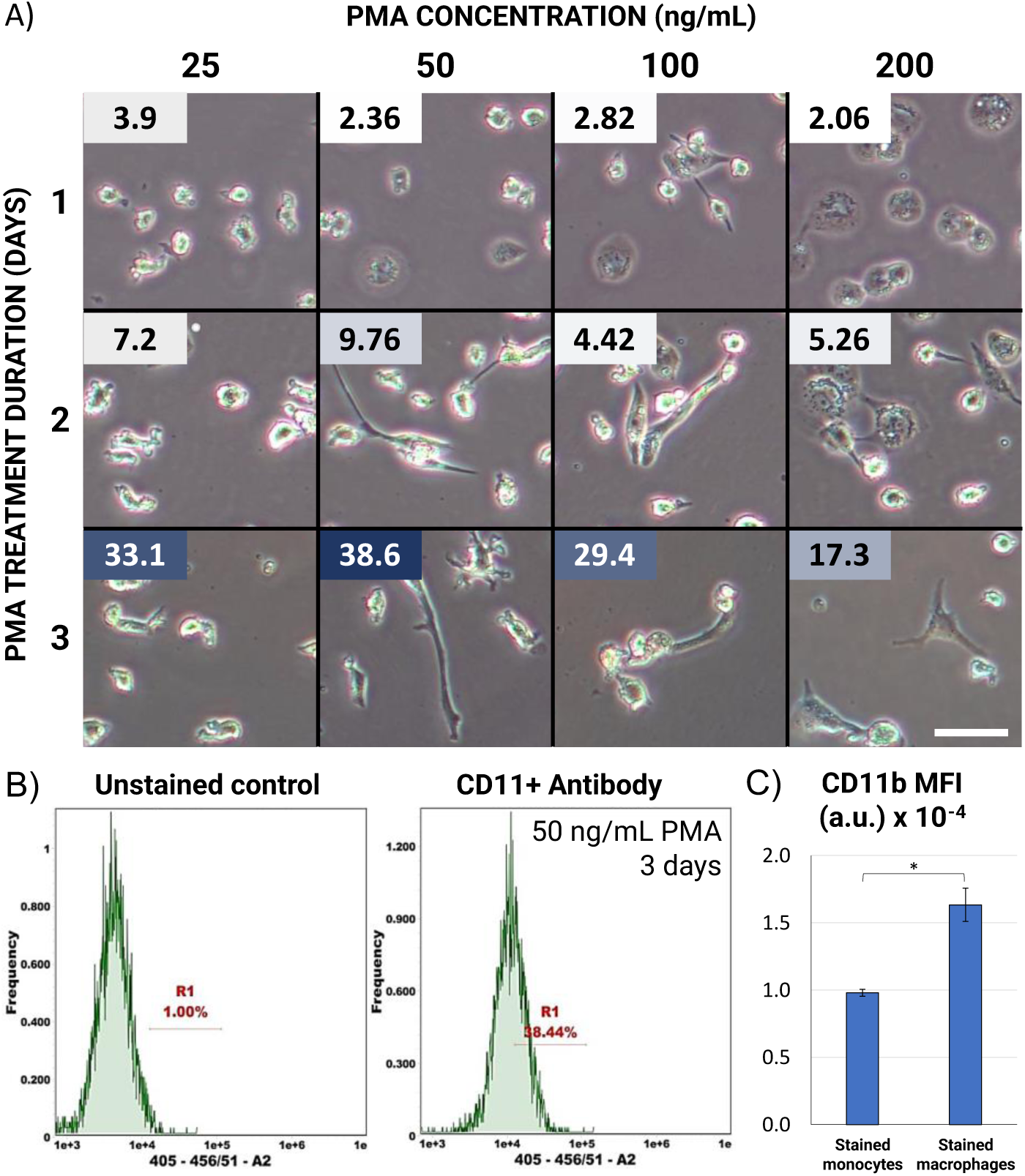
Optimization of PMA treatment for THP-1 monocyte differentiation. A) Morphology of THP-1 cells after treatment with PMA in well plate. Boxed number indicates percentage of CD11b+ cells. Scale bar = 100 μm. B) Representative histograms of PMA-treated THP-1 cells stained with a fluorescently labelled CD11b antibody, and unstained control to account for background fluorescence. C) Mean Fluorescent Intensity (MFI) of stained THP-1 “monocytes” (untreated) and “macrophages” (PMA-treated, 50ng/mL for 3 days). * p-value < 0.05. Error bars represent SEM.

After optimizing THP-1 differentiation, we cocultured HCT116 colonocytes and THP-1 cells in our preciously developed model of colonocyte-microbiota interactions in colorectal cancer^30^. In this model, colorectal cancer spheroids and colonic microbiota are cultured in separated, continuously perfused compartments and communicate via secreted metabolites through a porous membrane (**Figure 2A-C**). To include macrophages in this model, a suspension of THP-1 monocytes and HCT116 colonocytes in Matrigel was injected in the mammalian cell culture chamber on day 0 and perfused with media containing the optimized PMA concentration for 3 days to foster monocyte polarization during colonocyte spheroid formation before coculture with microbiota and/or treatment with inulin (**Figure 2D)**. Treatment of THP-1 monocytes in the device with PMA resulted in a significant increase in the expression of CD11b, with 60% of cells becoming positive and a 2.5-fold increase in MFI compared to untreated THP-1 monocytes in flask cultures (**Figure 2E, F**). These results confirmed the differentiation of THP-1 cells into a macrophage-like phenotype in our coculture model.

**Figure 2.**
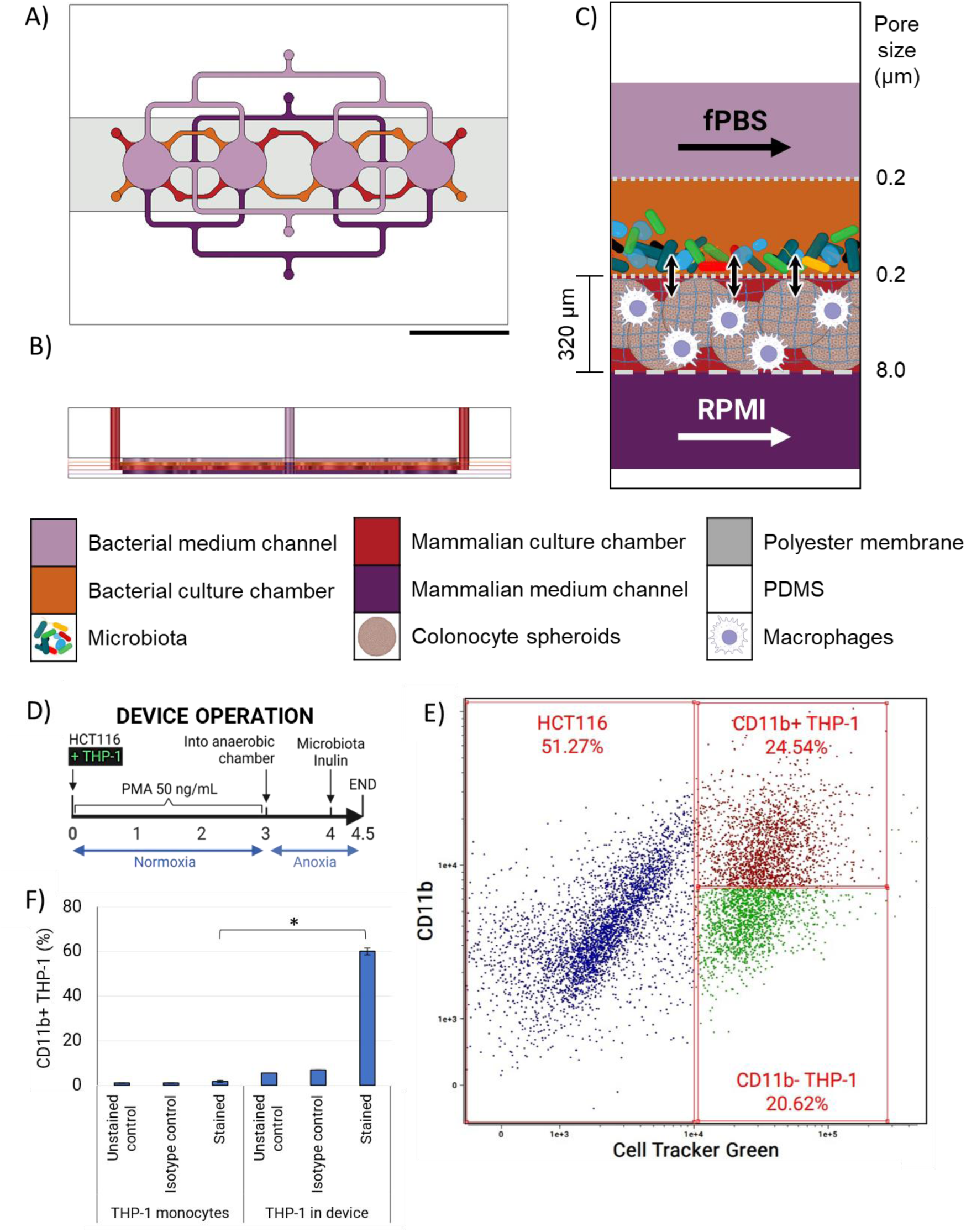
Microfluidic device design and operation. A) Top and B) cross-sectional views highlighting media channels and culture chambers. Scale bar = 1 cm. C) Schematic representation of device cocultures. Confined bacterial, colonocyte and macrophage populations interact via small molecules during perfusion with growth media. D) Experimental schedule of device coculture. E) Dot-plot of HCT116 – THP-1 cells after device coculture. THP-1 cells were loaded with CellTracker Green dye prior to coculture. The CD11b positivity threshold was defined based on unstained control. F) Percentage of CD11b+ THP-1 monocytes (control) and THP-1 cells after treatment with PMA during device coculture. * p-value < 0.05. Error bars represent SEM.

**Figure 2a.**
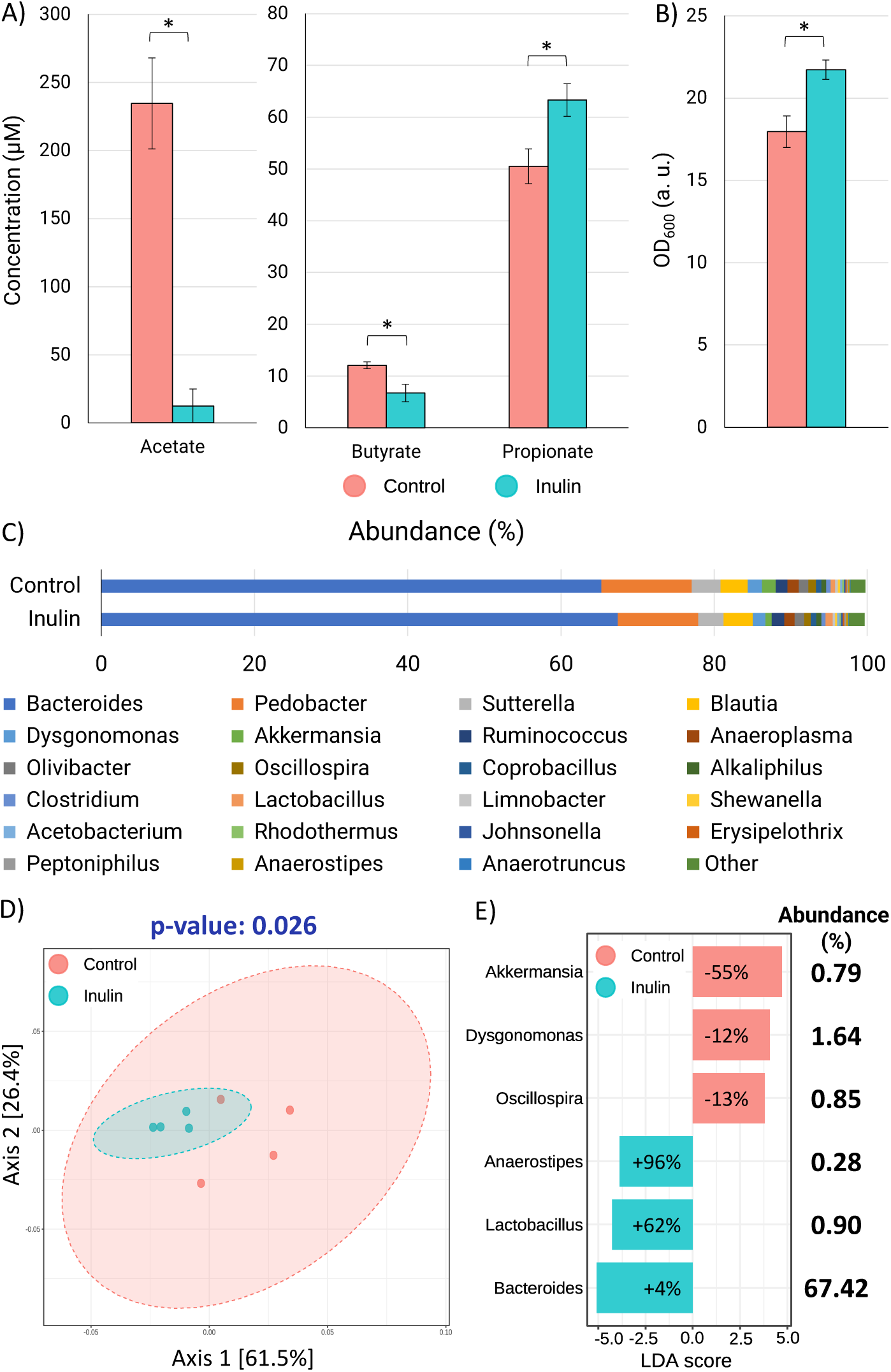
Effect of inulin on microbiota function, abundance, and composition. A) SCFA concentration in the bacterial culture chamber upon treatment with fPBS supplemented with 2.5% (w/v) inulin. B) Optical density of microbiota extracted from the device after inulin treatment. * p-value < 0.05. C) Microbiota composition at the genus level, D) β-diversity analysis, and E) LEfSe comparison of microbiota after treatment with inulin (2.5% w/v in fPBS) or control (fPBS) on chip. Percentages in the bars in E) correspond to relative change in the abundance of a genera normalized to abundance in control, and Abundance (%) corresponds to the composition in the inulin-treated microbiota.

### Inulin induces changes in microbiota abundance and function

To validate the use of our model to study the impact of dietary fiber on microbiota, we characterized the effect of inulin treatment on SCFA production and alterations in microbiota composition. Three SCFAs (acetate, propionate, and butyrate) were detected in the microbiota samples using GC-MS. Treatment with inulin resulted in a significant 95% (from 234 µM to 12 µM) decrease in acetate and a less pronounced decrease (44% (from 12 µM to 7 µM)) in butyrate levels, relative to the untreated controls. On the other hand, a significant 25% increase (from 50 µM to 63 µM) in propionate concentration was observed (**Figure 3A**), indicating differential effect of inulin exposure on the microbial community. Inulin treatment also induced bacterial proliferation, as noticed by a 21% increase (from 18 to 22) in optical density (**Figure 3B**).

**Figure 3.**
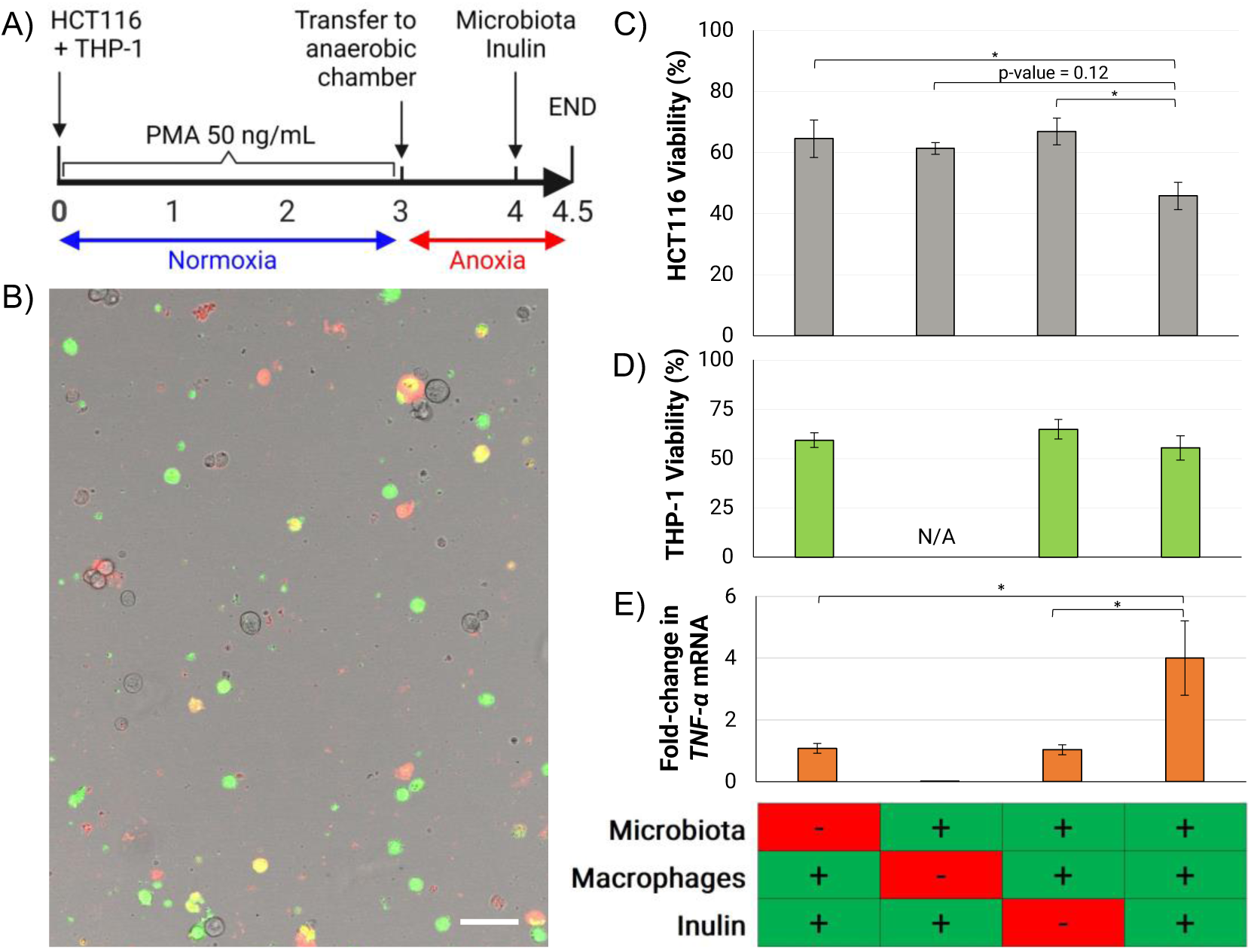
Effect of macrophages, inulin, and microbiota on colonocyte viability. A) Experimental schedule. B) Representative image of multicolor strategy to distinguish viability by cell type. Unstained: viable HCT116 cells. Red: dead HCT116 cells. Green: viable THP-1 cells. Yellow: dead THP-1 cells (Green + Red). Scale bar = 50 μm. C) HCT116 and D) THP-1 viability upon combinatorial treatment with microbiota and inulin. E) Associated fold-change in *TNF-α* transcription. * p-value < 0.05. Error bars represent SEM.

Metagenomic analysis revealed minimal changes in the composition of the community upon treatment with inulin. Regardless of inulin treatment, the microbiota cultured on chip was dominated by members of the *Bacteroides* genus (70%), and 24 other genera commonly associated with gastrointestinal microbiota, such as *Blautia*, *Ruminococcus*, and *Akkermansia*, that were present at an abundance higher than 1% (**Figure 3C**). The β-diversity analysis shows that while inulin treatment resulted in a significant change in overall microbiota composition, there was an overlap between ellipses that define a 95% confidence interval around the centroid of each treatment (**Figure 3D**). This result is consistent with the small but significant change in the abundance of members of the community upon treatment with inulin, including *Akkermansia* (0.45-fold), *Dysgonomonas* (0.88-fold), *Oscillospira* (0.87-fold), *Anaerostipes* (1.96-fold), and *Lactobacillus* (1.62-fold) (**Figure 3E**), as well as a 1.2-fold increase in the relative abundance of 6 genera (*Pseudobutyrivibrio, Anaerotruncus, Anaerobranca, Erysipelothrix, Turicibacter, Lachnospira, Peptoniphilus*) that did not reach statistical significance. The largest effect of inulin on microbiota composition was a 4% increase in the abundance of the genus *Bacteroidetes*, while the largest change in relative composition was a 1.96-fold increase in the abundance of the genus *Anaerostipes*.

### Inulin enhances macrophage-mediated colonocyte death in a microbiota-dependent manner

To evaluate the impact of inulin on macrophage-microbiota-colonocyte interaction, we cocultured HCT116 colonocytes with THP-1 macrophages and microbiota for 12 hours (**Figure 4A**). HCT116 spheroids without THP-1 macrophages, HCT116 spheroids and microbiota without THP-1 macrophages, and HCT-116, THP-1, and microbiota without inulin were used as controls to assess the impact of each component on HCT116 cell viability. The viability of each cell type was assessed employing a combination of CellTracker Green dye for labelling THP-1 cells and PI staining for labelling dead cells (**Figure 4B**). Inulin treatment of microbiota in coculture with HCT116 colonocytes and THP-1 macrophages resulted in a decrease of 17% in colonocyte viability relative to control treatments without macrophages, microbiota, or inulin (**Figure 4C**). In contrast, THP-1 viability remained relatively unchanged, regardless of the presence of microbiota or treatment with inulin (**Figure 4D**). The reduction in colonocyte viability correlated with a 4-fold increase in *TNF-α* transcription by THP-1 macrophages with respect to cocultures without either microbiota or inulin (**Figure 4E**), suggesting an effect of inulin metabolism by the microbiota on macrophage-colonocyte interaction. Treatment of macrophages with the SCFA mixture quantified in cocultures with microbiota and inulin (Inulin +) resulted in a 1.5-fold increase in *TNF-α* transcription compared to treatment with the SCFA mixture in cocultures without inulin (Inulin-) (**Figure 4**).

**Figure 4.**
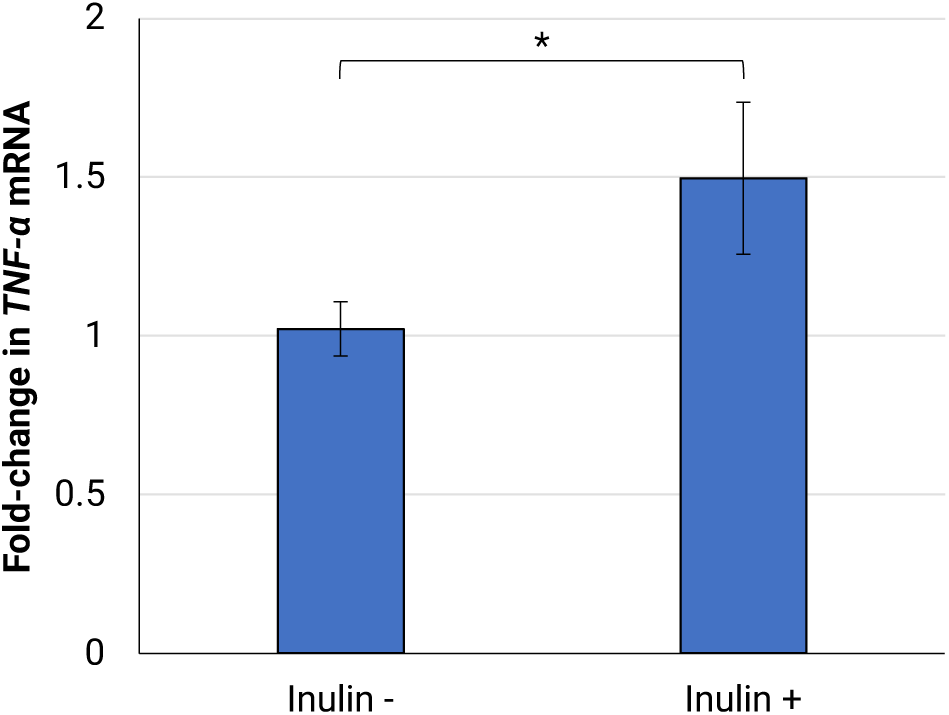
Effect of SCFA on *TNF-α* transcription in THP-1 macrophages. * p-value < 0.10, n = 7. Error bars represent SEM.

## Discussion

Dietary fiber intake significantly correlates with decreased incidence of CRC prospective human cohort studies^1,34^. Administration of inulin to murine CRC models has been shown to result in increased microbial abundance, changes in microbiota composition, and alterations in SCFA production^2,35^. In our model, inulin treatment significantly increased microbial proliferation quantified as optical density (**Figure 3B**), which agrees with the reported increased in cecal weight in mice and rats that has been partly attributed to increased bacterial abundance^36,37^. Inulin treatment also statistically altered the overall composition of the microbiota, although the impact on the abundance of most genera was not significant (**Figure 3C-D**). Members from the genus *Bacteroides*, an abundant member of human and murine intestinal microbiota that outcompetes other inulin fermenters such as bifidobacteria *in vitro*^38^, were detected in the microbiota cultured in our model and increased in abundance with inulin treatment. Therefore, the small impact of inulin on the overall microbiota composition could be a consequence of interspecies competition dominated by *Bacteroidetes* (**Figure 3E**). The abundance of the genera *Lactobacillus* and *Anaerostipes*, reported to increase in individuals upon ingestion of inulin^39–42^, also significantly increased in our model.

Inulin treatment resulted in a significant increase in propionate (**Figure 3A**), which is the most consistently reported effect of inulin administration in murine models^43–48^. The increase in propionate concentration also agrees with the high abundance of *Bacteroides* in the community, as members of this genus ferment inulin into propionate^49,50^. The effect of inulin administration on the levels of acetate in the colon *in vivo* have been inconsistent, with at least two studies showing either no effect or a decrease in abundance^43,45^. In our model, inulin treatment resulted in a significant decrease in extracellular acetate, which might have been caused by acetate conversion into acetyl-CoA for fatty acid biosynthesis and tricarboxylic acid synthesis under low oxygen conditions to support proliferation^51^. While inulin consumption is frequently associated with increased butyrate levels in mice and rats^44–46,48^, a decrease in butyrate was observed in our model. Crucially, inulin and fructose fermenters are known to produce acetate that is then employed by other species to produce butyrate^52^; therefore, the lack of an increase in butyrate could simply be a consequence of the short treatment time (12 hours) compared to weeks of inulin feeding in animal experiments^43–48^. It is also important to highlight that the response of gastrointestinal microbiota to dietary fiber interventions *in vivo* is heavily dependent on initial microbiota composition, which has resulted in significant inter-subject and inter-study variability^53–56^ and may explain the differences between some our results and other studies. Overall, our model recapitulated the increase in microbiota abundance and changes in SCFA production induced by inulin *in vivo*, which supports its use to study the effect of this dietary fiber on colonic microbiota *in vitro*.

Based on the immunomodulatory activity of the microbiota in the colon and the pivotal role of macrophages in CRC, we hypothesized that the changes in microbiota induced by inulin impact macrophage-colonocyte interaction in our coculture model. Our results show a macrophage-dependent increase in colonocyte death upon coculture with microbiota treated with inulin (**Figure 4C**). This correlated with an increase in the transcription of *TNF-α* (**Figure 4E**), a proinflammatory cytokine that of triggers apoptosis and necrosis in tumor-derived cell lines^21^, including CRC cell lines^57,58^. The increase in *TNF-α* transcription correlated with a decrease in acetate and an increase in propionate with inulin (**Figure 3A**), which agrees with the strong inhibition of TNF-α production in myeloid cells by acetate^59^. Treatment of THP-1 macrophages with a mixture of SCFA matching the concentrations in device cocultures with inulin partially recapitulated the increase in *TNF-α* transcription observed in device cocultures (**Figure 4**). Inulin-induced changes in the abundance of other microbial metabolites that affect TNF-α production in macrophages, such as lipopolysaccharide and indole derivatives^60^, may also explain the observed effect of inulin on macrophage activity, and require further characterization.

A strong association between dietary fiber consumption and lower CRC incidence reported by metanalyses of prospective cohort studies^1,34^. This preventive activity of dietary fiber against colorectal cancer has been attributed to the increased production of total SCFA by microbiota and its direct proapoptotic effect on colonocytes^61,62^. Importantly, a large number of dietary intervention studies in humans have found little to no effect of inulin ingestion on fecal SCFA concentration^63–68^. Our results suggest that dietary fiber may induce colonocyte death via changes in macrophage activity even in the context of a decreased total abundance of SCFA. The microbiota-mediated immunomodulatory activity by inulin proposed here is similar to the improved Natural Killer cell cytotoxicity in a rat model of CRC upon inulin administration ^69^, as well as the reduction in xenograft tumor growth in mice upon inulin administration via microbiota-dependent T-cell activation^70^. Since both diet and immune cell activity are key factors in CRC, the mechanisms underlying the potential immunomodulatory effect of dietary fiber on the immune component of the tumor microenvironment require further study.

The role of macrophages in CRC is controversial^71,72^. While *in vitro* studies have shown that macrophages induce CRC colonocyte proliferation and migration^16–20^, in epidemiological studies show that high macrophage infiltration in CRC tumors is often associated with better prognosis^73^. Importantly, *in vitro* studies have failed to consider the hypoxic, ECM-rich, three-dimensional microenvironment of tumor tissue^13^, as well as the impact of the microbiota and its products on macrophage activity. Our results suggest that macrophages may display cytotoxic activity against colonocytes *in vitro*, more consistent with epidemiological data, when cultured in a physiologically relevant microenvironment that contains microbiota and dietary molecules. These observations support the potential of microfluidic technology to study complex host-microbiota interactions *in vitro* in a manner that is more relevant than conventional cell culture experiments^74^.

While the changes in microbiota composition upon inulin treatment we observed are in line with reported effects of inulin *in vivo*, a more extended coculture time might result in larger changes in composition which would better recapitulate the shifts in composition observed *in vivo.* The mechanisms of inulin-microbiota-macrophage interaction proposed here are based on correlation among variables and therefore require further investigation to demonstrate causality. This could be accomplished by employing selective TNFR1 inhibitors^75^, synthetic microbial communities^56^, and macrophages with diminished TNF-α production capacity^76^. Further characterization of the microbiota via metabolomics, as well as macrophages via RNA sequencing and expanded immunotyping, would be useful to identify correlations between microbial metabolism, macrophage activation, and colonocyte viability. The use of more relevant sources of macrophages, such as bone-marrow derived monocytes, would also increase the biological relevance of future studies^77^. Finally, a diverse group of dietary fibers and more extended experimental times may increase our understanding of the dynamics of dietary fiber fermentation and changes in microbiota and macrophage activity.

## Methods

### Mammalian cell culture and reagents

The human colorectal cancer cell line HCT116 (CCL-247) was obtained from ATCC and cultured in RPMI 1640 medium (Corning) supplemented with 10% FBS (Atlanta Biologicals), GlutaMAX, HEPES, and NEAA (Gibco). The human monocyte cell line THP-1 was obtained from ATCC and cultured in RPMI 1640 medium (Corning) supplemented with 10% FBS (Atlanta Biologicals), 1% GlutaMAX, 1% HEPES, 1% NEAA (Gibco), and 2-mercaptoethanol (Sigma-Aldrich) to a final concentration of 0.05 mM. THP-1 cultures were maintained at a density between 2×10^5^ and 9×10^5^ cells/mL. For device coculture experiments, THP-1 cells were labelled with 2 µM CellTracker Green CMFDA (ThermoFischer Scientific) in serum-free media for 45 minutes.

### Optimization of THP-1 differentiation

THP-1 monocytes were differentiated to macrophages with phorbol 12-myristate 13-acetate (PMA, Sigma-Aldrich). THP-1 differentiation was optimized using ∼5×10^5^ cells/mL in a 1 mL culture tube and treated with PMA to a final concentration of 25, 50, 100 or 200 ng/mL. Cells were incubated at 37°C for 1, 2, or 3 days. To detach PMA-treated THP-1 cells from culture tubes for flow cytometry analysis, the PMA-containing media was replaced with PBS with 10 mM EDTA, incubated on ice for 15 minutes, and detached by repeated pipetting.

### Microfluidic device construction and operation

Device construction and operation was performed as previously described^30^. The device consists of four PDMS layers separated by three porous polyester membranes (**Figure**). Thin patterned PDMS layers were obtained by pouring uncured PDMS mix (Sylgard 184®) on a 3D-printed patterned mold (Stratasys, Inc.) with a total height of 500 μm and a pattern height of 160 μm, and then the PDMS was cured for 1 hour at 70 °C. Biopsy punches were used to create culture chambers (D = 5 mm, final chamber volume of ∼10 μL) and open access to the perfusion channels and culture chambers (D = 1 mm). The device was assembled layer-by-layer using a thin layer of uncured PDMS as glue, and a polyester membrane was sandwiched between each pair of layers. The device was connected to media reservoirs by flexible 23-Gauge Tygon® medical tubing (Saint Gobain) using 20-Gauge stainless steel connectors detached from dispensing needles (Jensen Tools). To minimize the formation of bubbles during operation, the assembled device was placed underwater and vacuumed to a final pressure of 3×10^-2^ mbar for 24 hours; then, the device was autoclaved (121 C, 16 PSIG, 45 minutes) and kept covered in sterile water during operation. This protocol sterilized the device and prevented bubble formation. The device was held underwater for the duration of the experiment and only brought out of water for cell injection.

For colonocyte injection, themammalian culture chambers in the device were seeded with 50% v/v Matrigel diluted in RPMI medium containing either ∼4×10^6^ HCT116 cells/mL or ∼2×10^6^ HCT116 and ∼2×10^6^ labelled THP-1 cells/mL. After cell seeding, devices were perfused with antibiotic-free RPMI through the mammalian medium channel, while PBS supplemented with 50 ng/mL of PMA was perfused through the bacterial chamber. On day 4 of culture, the bacterial chamber media was replaced with PMA-free PBS, and devices were transferred and operated in an anaerobic chamber for 24 hours. On day 5, murine fecal microbiota was obtained from freshly-voided fecal pellets obtained from 8 to 12 weeks-old wild-type C57BL/6 female mice fed a standard chow diet and processed as previously described to isolate microbiota and prepare fecal PBS (fPBS)^30^. The fecal slurry containing microbiota was introduced into the bacterial chamber, and the PBS in the bacterial medium reservoir was replaced with sterile-filtered fPBS or fPBS supplemented with inulin at a concentration of 2.5% (w/v) (Spectrum Chemical). Cocultures proceeded for 12 hours.

For sample collection, the microbiota in the bacterial culture chamber was collected by pipetting, to a final volume of ∼10 µL per chamber. Bacteria were pelleted by centrifugation at 10000g for 10 minutes. The supernatant was used for SCFA analysis by GC-MS while the bacterial pellet was resuspended in 500 µL of PBS, the OD_600_ noted, and centrifuged (10000g, 10 minutes) to obtain a bacterial cell pellet. All pellets were stored at −80 °C until further processing. Next, the mammalian hydrogels were recovered by carefully disassembling the device with a scalpel. For flow cytometry and viability analysis, the mammalian hydrogels were placed on ice in PBS with 10 mM EDTA for 15 minutes and disaggregated by repeated pipetting to obtain single cell suspensions.

### Immunostaining and flow cytometry

For immunostaining, Fc blocking was performed using human IgG (Sigma-Aldrich) at a concentration of 4 µg/10^6^ cells for 15 minutes at room temperature. Cells were stained with Human CD11b/Integrin alpha M Alexa Fluor® 405-conjugated Antibody (R&D Systems) at a concentration of 1 µg/10^6^ cells according to manufacturer’s protocols, using normal mouse IgG2b Alexa Fluor 405 (Santa Cruz Biotechnology) as isotype control. Flow cytometry was performed using a CellStream benchtop flow cytometer (Millipore Sigma). Single color controls were used to create a compensation matrix for signal bleed between fluorophores. Single cells were acquired using a 0.6-1 Aspect Ratio as the criterion. Dead cells were excluded with Propidium Iodide (PI) staining (1.5 μM for 5 minutes).

### Mammalian cell viability evaluation

Single cell suspensions were stained with 1.5 μM PI and incubated for 15 minutes in the dark at room temperature. Viability was evaluated using a Leica TCS SP5 confocal microscope. At least 100 cells were counted per sample, and 4 hydrogels were processed per treatment.

### SCFA analysis

SCFA quantification was carried out by the Integrated Metabolomics Analysis Core at Texas A&M University. Metabolites were extracted from samples using ethyl acetate and the levels of 6 SCFAs (acetic, butyric, isobutyric, isovaleric, propionic, and valeric acid) were quantified using GC-MS.

Isotopically labelled n-butyrate was used as the internal control and was spiked into all samples prior to extraction. Samples were diluted 10-fold in PBS before extraction. SCFAs were detected and quantified on a gas chromatography triple quadrupole mass spectrometer (TSQ EVO 8000, Thermo Scientific, Waltham, MA). Chromatographic separation was achieved on a ZB WAX Plus, 30 m x 0.25mm x 0.25 µm column (Phenomenex). The MS data and retention times were acquired in full scan mode from m/z 40-500 for the individual target compounds. The injector, MS transfer line and ion source were maintained at 230 °C, 240 °C and 240 °C respectively. The flow rate of helium carrier gas was kept at 1 mL/min. Samples were maintained at room temperature on an autosampler before injection. 1 µl of the extracted sample was injected with a split ratio of 20:1. The ionization was carried out in the electron impact (EI) mode at 70 eV. Sample acquisition and analysis was performed with TraceFinder 3.3 (Thermo Scientific).

### Microbiota composition analysis

DNA from bacterial communities was extracted by using the DNeasy PowerSoil Kit (QIAGEN) according to manufacturer’s instructions, and the v4 region of the 16S rRNA gene was sequenced using the MiSeq Illumina Platform (FERA Diagnostics and Biologicals). Bioinformatic analysis was performed using Microbiome Analyst (www.microbiomeanalyst.ca) at the genus level. Features with singlets were removed prior to analysis. Features with single readings were removed before analysis. For comparative analysis, only features with a read count higher than 4 in 50% of samples were included, and 10% of features with the lowest coefficient of variation were removed to ameliorate data sparsity and improve statistical power^78^. Data was scaled using Cumulative Sum Scaling. The Bradis-Curtis Index distance metric was used with PERMANOVA as the statistical method. LEfSe analysis was performed using LDA > 2.0 as the significance filter and p < 0.05 was used for statistical significance.

### Effect of SCFA on TNF-α transcription

To evaluate the effect of SCFA on *TNF-α* transcription, 5×10^5^ THP-1 cells in 1 mL of RPMI were seeded into a 12-well culture plate and treated with 50 ng/mL PMA for 72 hours. PMA-containing media was then replaced with fresh RPMI, and cells were incubated for 24 hours. Cells were treated with RPMI medium supplemented with SCFA for 12 hours according to **Table**.

**Table 1.** Composition of SCFA mixtures (µM).

| SCFA | Inulin - | Inulin + |
| --- | --- | --- |
| Sodium Acetate | 234.6 | 12.5 |
| Sodium Butyrate | 12.1 | 6.7 |
| Sodium Propionate | 50.5 | 63.3 |

### Gene expression analysis

RNA was extracted from mammalian cells using the RNeasy Mini Kit (QIAGEN) following the manufacturer’s instructions. Genomic DNA in the extracted RNA was eliminated by digesting with DNAse (QIAGEN). cDNA synthesis was performed using qScript™ cDNA SuperMix (QuantaBio) using 100 ng of RNA sample in a 10 µL reaction. Quantitative PCR was carried out in a Lightcycler® 96 (Roche) using FastStart Universal SYBR Green Master (Roche). Primers were designed using Primer Blast (NCBI), and amplicon size and specificity were confirmed by melting peak analysis and agarose gel electrophoresis of the reaction products. Each reaction mix contained 1/40^th^ of the cDNA pool obtained per sample and a total primer concentration of 400 nM. The PCR regime consisted of preincubation for 10 minutes at 95 °C and 45 amplification cycles (95 °C x 15 s, 65 °C x 30 s, 72 °C x 45 s). Data were processed using the 2^-ΔΔCt^ method. Multiple genes were evaluated as endogenous qPCR controls, including *18s*, *YWHAZ*, *PMM1*, *UBC*, *IPO8*, and *VPS29*; from these genes, *PMM1* showed the most stable expression level and was employed as endogenous control. Sequences for all used primers are provided in **Table 2**.

**Table 2.**
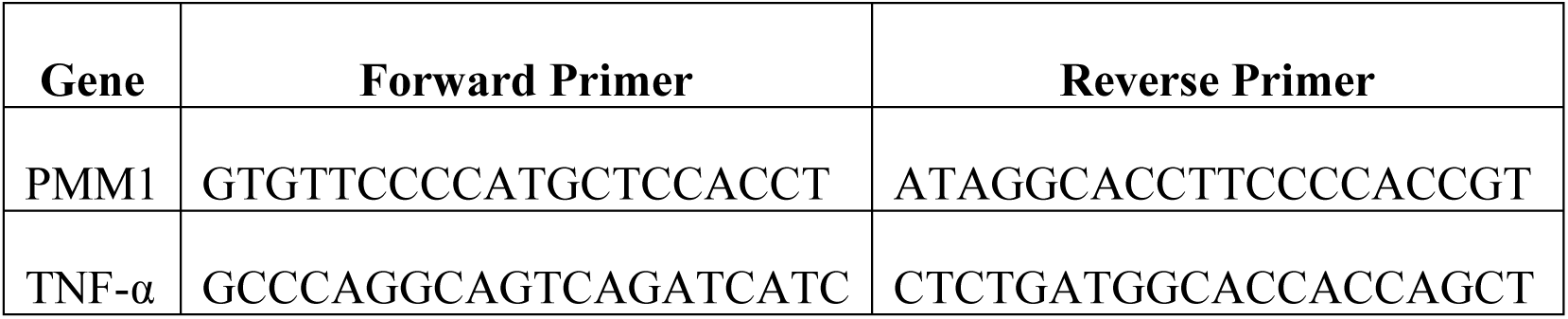
Primer sequences for gene expression analysis.

### Statistical analysis

For testing statistical significance, unpaired Student’s t-tests were performed on sets of data with two experimental conditions. One-way ANOVAs were used for comparisons among multiple experimental conditions and during RTqPCR data analysis. For RTqPCR data analysis, significance in gene expression changes was determined by comparing ΔCt values across treatments, as gene expression data is log normally distributed^79^. The assumption of equality of variances among data sets was confirmed by using the Levene’s test, and normality was validated using the Shapiro-Wilk test. All experiments were performed at least in duplicate, and coculture experiments were performed in triplicate.

## Author contributions

D. P., M. M., and A. J. designed the research. D. P. and M. M. performed the experiments. D. P., M. M., S. C., A. H. and A. J. analyzed the data. D. P., M. M., and A. J. wrote the article with input from A. H and S. C. All authors reviewed, discussed, and edited the manuscript.

## Acknowledgements

This work was partially supported by funds from the Ray B. Nesbitt Chair endowment to A. J.

## References

1. Veettil, S. K. et al. Role of Diet in Colorectal Cancer Incidence: Umbrella Review of Meta-analyses of Prospective Observational Studies. JAMA Netw. Open 4, e2037341 (2021).

2. Pool-Zobel, B. L. Inulin-type fructans and reduction in colon cancer risk: review of experimental and human data. Br. J. Nutr. 93 Suppl 1, S73–90 (2005).

3. So, D. et al. Dietary fiber intervention on gut microbiota composition in healthy adults: a systematic review and meta-analysis. Am. J. Clin. Nutr. 107, 965–983 (2018).

4. Bishehsari, F. et al. Dietary Fiber Treatment Corrects the Composition of Gut Microbiota, Promotes SCFA Production, and Suppresses Colon Carcinogenesis. Genes 9, (2018).

5. Makki, K., Deehan, E. C., Walter, J. & Bäckhed, F. The Impact of Dietary Fiber on Gut Microbiota in Host Health and Disease. Cell Host Microbe 23, 705–715 (2018).

6. Wu, X. et al. Effects of the intestinal microbial metabolite butyrate on the development of colorectal cancer. J. Cancer 9, 2510–2517 (2018).

7. Waldecker, M., Kautenburger, T., Daumann, H., Busch, C. & Schrenk, D. Inhibition of histone-deacetylase activity by short-chain fatty acids and some polyphenol metabolites formed in the colon. J. Nutr. Biochem. 19, 587–593 (2008).

8. Sahuri-Arisoylu, M. et al. Acetate Induces Growth Arrest in Colon Cancer Cells Through Modulation of Mitochondrial Function. Front. Nutr. 8, 588466 (2021).

9. Oliveira, C. S. F. et al. Cathepsin D protects colorectal cancer cells from acetate-induced apoptosis through autophagy-independent degradation of damaged mitochondria. Cell Death Dis. 6, e1788–e1788 (2015).

10. Marques, C. et al. Acetate-induced apoptosis in colorectal carcinoma cells involves lysosomal membrane permeabilization and cathepsin D release. Cell Death Dis. 4, e507–e507 (2013).

11. Ryu, T. Y. et al. Downregulation of PRMT1, a histone arginine methyltransferase, by sodium propionate induces cell apoptosis in colon cancer. Oncol. Rep. 41, 1691–1699 (2019).

12. Liong, M.-T. Roles of Probiotics and Prebiotics in Colon Cancer Prevention: Postulated Mechanisms and In-vivo Evidence. Int. J. Mol. Sci. 9, 854–863 (2008).

13. Li, J., Chen, D. & Shen, M. Tumor Microenvironment Shapes Colorectal Cancer Progression, Metastasis, and Treatment Responses. Front. Med. 9, (2022).

14. van Ravenswaay Claasen, H. H., Kluin, P. M. & Fleuren, G. J. Tumor infiltrating cells in human cancer. On the possible role of CD16+ macrophages in antitumor cytotoxicity. Lab. Investig. J. Tech. Methods Pathol. 67, 166–174 (1992).

15. McBride, W. H. Phenotype and functions of intratumoral macrophages. Biochim. Biophys. Acta BBA - Rev. Cancer 865, 27–41 (1986).

16. Luput, L. et al. Tumor-associated macrophages favor C26 murine colon carcinoma cell proliferation in an oxidative stress-dependent manner. Oncol. Rep. 37, 2472–2480 (2017).

17. Miao, H. et al. Macrophage ABHD5 promotes colorectal cancer growth by suppressing spermidine production by SRM. Nat. Commun. 7, 11716 (2016).

18. Lan, J. et al. M2 Macrophage-Derived Exosomes Promote Cell Migration and Invasion in Colon Cancer. Cancer Res. 79, 146–158 (2019).

19. Jedinak, A., Dudhgaonkar, S. & Sliva, D. Activated macrophages induce metastatic behavior of colon cancer cells. Immunobiology 215, 242–249 (2010).

20. Vinnakota, K. et al. M2-like macrophages induce colon cancer cell invasion via matrix metalloproteinases. J. Cell. Physiol. 232, 3468–3480 (2017).

21. Wang, X. & Lin, Y. Tumor necrosis factor and cancer, buddies or foes? Acta Pharmacol. Sin. 29, 1275–1288 (2008).

22. de Looff, M., de Jong, S. & Kruyt, F. A. E. Multiple Interactions Between Cancer Cells and the Tumor Microenvironment Modulate TRAIL Signaling: Implications for TRAIL Receptor Targeted Therapy. Front. Immunol. 10, 1530 (2019).

23. Baay, M., Brouwer, A., Pauwels, P., Peeters, M. & Lardon, F. Tumor Cells and Tumor-Associated Macrophages: Secreted Proteins as Potential Targets for Therapy. Clin. Dev. Immunol. 2011, 565187 (2011).

24. Han, C., Zhang, C., Wang, H. & Zhao, L. Exosome-mediated communication between tumor cells and tumor-associated macrophages: implications for tumor microenvironment. Oncoimmunology 10, 1887552 (2021).

25. Chen, Q., Nair, S. & Ruedl, C. Microbiota regulates the turnover kinetics of gut macrophages in health and inflammation. Life Sci. Alliance 5, (2022).

26. Mola, S., Pandolfo, C., Sica, A. & Porta, C. The Macrophages-Microbiota Interplay in Colorectal Cancer (CRC)-Related Inflammation: Prognostic and Therapeutic Significance. Int. J. Mol. Sci. 21, 6866 (2020).

27. Fang, H. et al. Lipopolysaccharide-induced macrophage inflammatory response is regulated by SHIP. J. Immunol. Baltim. Md 1950 173, 360–366 (2004).

28. Ji, J. et al. Microbial metabolite butyrate facilitates M2 macrophage polarization and function. Sci. Rep. 6, 1–10 (2016).

29. Schulthess, J. et al. The Short Chain Fatty Acid Butyrate Imprints an Antimicrobial Program in Macrophages. Immunity 50, 432–445.e7 (2019).

30. Penarete-Acosta, D. et al. A microfluidic co-culture model for investigating colonocytes– microbiota interactions in colorectal cancer. Lab. Chip 24, 3690–3703 (2024).

31. Park, E. K. et al. Optimized THP-1 differentiation is required for the detection of responses to weak stimuli. Inflamm. Res. Off. J. Eur. Histamine Res. Soc. Al 56, 45–50 (2007).

32. Starr, T., Bauler, T. J., Malik-Kale, P. & Steele-Mortimer, O. The phorbol 12-myristate-13-acetate differentiation protocol is critical to the interaction of THP-1 macrophages with Salmonella Typhimurium. PLOS ONE 13, e0193601 (2018).

33. Lund, M. E., To, J., O’Brien, B. A. & Donnelly, S. The choice of phorbol 12-myristate 13-acetate differentiation protocol influences the response of THP-1 macrophages to a pro-inflammatory stimulus. J. Immunol. Methods 430, 64–70 (2016).

34. Ma, Y. et al. Dietary fiber intake and risks of proximal and distal colon cancers: A meta-analysis. Medicine (Baltimore) 97, e11678 (2018).

35. Pool-Zobel, B. L. & Sauer, J. Overview of Experimental Data on Reduction of Colorectal Cancer Risk by Inulin-Type Fructans. J. Nutr. 137, 2580S–2584S (2007).

36. Cummings, J. H., Macfarlane, G. T. & Englyst, H. N. Prebiotic digestion and fermentation. Am. J. Clin. Nutr. 73, 415s–420s (2001).

37. Xiao, J., Metzler-Zebeli, B. U. & Zebeli, Q. Gut Function-Enhancing Properties and Metabolic Effects of Dietary Indigestible Sugars in Rodents and Rabbits. Nutrients 7, 8348–8365 (2015).

38. Falony, G., Calmeyn, T., Leroy, F. & De Vuyst, L. Coculture Fermentations of Bifidobacterium Species and Bacteroides thetaiotaomicron Reveal a Mechanistic Insight into the Prebiotic Effect of Inulin-Type Fructans. Appl. Environ. Microbiol. 75, 2312–2319 (2009).

39. Hiel, S. et al. Link between gut microbiota and health outcomes in inulin-treated obese patients: Lessons from the Food4Gut multicenter randomized placebo-controlled trial. Clin. Nutr. 39, 3618–3628 (2020).

40. Vandeputte, D. et al. Prebiotic inulin-type fructans induce specific changes in the human gut microbiota. Gut 66, 1968–1974 (2017).

41. Le Bastard, Q. et al. The effects of inulin on gut microbial composition: a systematic review of evidence from human studies. Eur. J. Clin. Microbiol. Infect. Dis. 39, 403–413 (2020).

42. Wang, X., Wang, T., Zhang, Q., Xu, L. & Xiao, X. Dietary Supplementation with Inulin Modulates the Gut Microbiota and Improves Insulin Sensitivity in Prediabetes. Int. J. Endocrinol. 2021, 5579369 (2021).

43. Fernández, J. et al. Traditional Processed Meat Products Re-designed Towards Inulin-rich Functional Foods Reduce Polyps in Two Colorectal Cancer Animal Models. Sci. Rep. 9, 14783 (2019).

44. Rao, C. V., Chou, D., Simi, B., Ku, H. & Reddy, B. S. Prevention of colonic aberrant crypt foci and modulation of large bowel microbial activity by dietary coffee fiber, inulin and pectin. Carcinogenesis 19, 1815–1819 (1998).

45. Poulsen, M., Mølck, A.-M. & Jacobsen, B. L. Different effects of short- and long-chained fructans on large intestinal physiology and carcinogen-induced aberrant crypt foci in rats. Nutr. Cancer 42, 194–205 (2002).

46. Femia, A. P. et al. Antitumorigenic activity of the prebiotic inulin enriched with oligofructose in combination with the probiotics Lactobacillus rhamnosus and Bifidobacterium lactis on azoxymethane-induced colon carcinogenesis in rats. Carcinogenesis 23, 1953–1960 (2002).

47. Wang, Z. et al. Inulin alleviates inflammation of alcoholic liver disease via SCFAs-inducing suppression of M1 and facilitation of M2 macrophages in mice. Int. Immunopharmacol. 78, 106062 (2019).

48. Nakajima, H. et al. Inulin reduces visceral adipose tissue mass and improves glucose tolerance through altering gut metabolites. Nutr. Metab. 19, 50 (2022).

49. Rios-Covian, D., Salazar, N., Gueimonde, M. & de los Reyes-Gavilan, C. G. Shaping the Metabolism of Intestinal Bacteroides Population through Diet to Improve Human Health. Front. Microbiol. 8, (2017).

50. Louis, P. & Flint, H. J. Formation of propionate and butyrate by the human colonic microbiota. Environ. Microbiol. 19, 29–41 (2017).

51. De Mets, F., Van Melderen, L. & Gottesman, S. Regulation of acetate metabolism and coordination with the TCA cycle via a processed small RNA. Proc. Natl. Acad. Sci. 116, 1043–1052 (2019).

52. Lee, H. L. et al. Targeted Approaches for In Situ Gut Microbiome Manipulation. Genes 9, 351 (2018).

53. Birkeland, E. et al. Prebiotic effect of inulin-type fructans on faecal microbiota and short-chain fatty acids in type 2 diabetes: a randomised controlled trial. Eur. J. Nutr. 59, 3325–3338 (2020).

54. Poeker, S. A. et al. Stepwise Development of an in vitro Continuous Fermentation Model for the Murine Caecal Microbiota. Front. Microbiol. 10, (2019).

55. Holmes, Z. C. et al. Microbiota responses to different prebiotics are conserved within individuals and associated with habitual fiber intake. Microbiome 10, 114 (2022).

56. Donohoe, D. R. et al. A Gnotobiotic Mouse Model Demonstrates that Dietary Fiber Protects Against Colorectal Tumorigenesis in a Microbiota- and Butyrate–Dependent Manner. Cancer Discov. 4, 1387–1397 (2014).

57. Fan, R., Naqvi, K., Patel, K., Sun, J. & Wan, J. Evaporation-based microfluidic production of oil-free cell-containing hydrogel particles. Biomicrofluidics 9, 052602 (2015).

58. Wilson, C. A. & Browning, J. L. Death of HT29 adenocarcinoma cells induced by TNF family receptor activation is caspase-independent and displays features of both apoptosis and necrosis. Cell Death Differ. 9, 1321–1333 (2002).

59. Eslick, S. et al. Weight Loss and Short-Chain Fatty Acids Reduce Systemic Inflammation in Monocytes and Adipose Tissue Macrophages from Obese Subjects. Nutrients 14, 765 (2022).

60. Krishnan, S. et al. Gut Microbiota-Derived Tryptophan Metabolites Modulate Inflammatory Response in Hepatocytes and Macrophages. Cell Rep. 23, 1099–1111 (2018).

61. O’Keefe, S. J. D. Diet, microorganisms and their metabolites, and colon cancer. Nat. Rev. Gastroenterol. Hepatol. 13, 691–706 (2016).

62. Fung, K. Y. C. et al. Butyrate-induced apoptosis in HCT116 colorectal cancer cells includes induction of a cell stress response. J. Proteome Res. 10, 1860–1869 (2011).

63. Ramirez-Farias, C. et al. Effect of inulin on the human gut microbiota: stimulation of Bifidobacterium adolescentis and Faecalibacterium prausnitzii. Br. J. Nutr. 101, 541–550 (2008).

64. Baxter, N. T. et al. Dynamics of Human Gut Microbiota and Short-Chain Fatty Acids in Response to Dietary Interventions with Three Fermentable Fibers. mBio 10, e02566–18 (2019).

65. Salazar, N. et al. Inulin-type fructans modulate intestinal Bifidobacterium species populations and decrease fecal short-chain fatty acids in obese women. Clin. Nutr. 34, 501–507 (2015).

66. Healey, G. et al. Habitual dietary fibre intake influences gut microbiota response to an inulin-type fructan prebiotic: a randomised, double-blind, placebo-controlled, cross-over, human intervention study. Br. J. Nutr. 119, 176–189 (2018).

67. Liu, F. et al. Fructooligosaccharide (FOS) and Galactooligosaccharide (GOS) Increase Bifidobacterium but Reduce Butyrate Producing Bacteria with Adverse Glycemic Metabolism in healthy young population. Sci. Rep. 7, 11789 (2017).

68. Holscher, H. D. et al. Agave Inulin Supplementation Affects the Fecal Microbiota of Healthy Adults Participating in a Randomized, Double-Blind, Placebo-Controlled, Crossover Trial. J. Nutr. 145, 2025–2032 (2015).

69. Roller, M., Pietro Femia, A., Caderni, G., Rechkemmer, G. & Watzl, B. Intestinal immunity of rats with colon cancer is modulated by oligofructose-enriched inulin combined with Lactobacillus rhamnosus and Bifidobacterium lactis. Br. J. Nutr. 92, 931–938 (2004).

70. Li, Y. et al. Prebiotic-Induced Anti-tumor Immunity Attenuates Tumor Growth. Cell Rep. 30, 1753–1766.e6 (2020).

71. Braster, R., Bögels, M., Beelen, R. H. J. & van Egmond, M. The delicate balance of macrophages in colorectal cancer; their role in tumour development and therapeutic potential. Immunobiology 222, 21–30 (2017).

72. Zhong, X., Chen, B. & Yang, Z. The Role of Tumor-Associated Macrophages in Colorectal Carcinoma Progression. Cell. Physiol. Biochem. 45, 356–365 (2018).

73. Li, J. et al. Tumor-associated macrophage infiltration and prognosis in colorectal cancer: systematic review and meta-analysis. Int. J. Colorectal Dis. 35, 1203–1210 (2020).

74. Bein, A. et al. Microfluidic Organ-on-a-Chip Models of Human Intestine. Cell. Mol. Gastroenterol. Hepatol. 5, 659–668 (2018).

75. Zhang, N., Wang, Z. & Zhao, Y. Selective inhibition of Tumor necrosis factor receptor-1 (TNFR1) for the treatment of autoimmune diseases. Cytokine Growth Factor Rev. 55, 80–85 (2020).

76. Covarrubias, S. et al. High-Throughput CRISPR Screening Identifies Genes Involved in Macrophage Viability and Inflammatory Pathways. Cell Rep. 33, 108541 (2020).

77. Tedesco, S. et al. Convenience versus Biological Significance: Are PMA-Differentiated THP-1 Cells a Reliable Substitute for Blood-Derived Macrophages When Studying in Vitro Polarization? Front. Pharmacol. 9, 71 (2018).

78. Dhariwal, A. et al. MicrobiomeAnalyst: a web-based tool for comprehensive statistical, visual and meta-analysis of microbiome data. Nucleic Acids Res. 45, W180–W188 (2017).

79. Derveaux, S., Vandesompele, J. & Hellemans, J. How to do successful gene expression analysis using real-time PCR. Methods 50, 227–230 (2010).

